# Structural basis for the simultaneous recognition of NEMO and acceptor ubiquitin by the HOIP NZF1 domain

**DOI:** 10.1101/2022.02.25.482043

**Authors:** Simin Rahighi, Mamta Iyer, Hamid Oveisi, Sammy Nasser, Vincent Duong

## Abstract

Ubiquitination of NEMO by the linear ubiquitin chain assembly complex (LUBAC) is essential for activating the canonical NF-κB signaling pathway. While the NZF1 domain of the HOIP subunit of LUBAC recognizes the NEMO substrate, it is unclear how it cooperates with the catalytic domains in the ubiquitination process. Here, we report a crystal structure of NEMO in complex with HOIP NZF1 and linear diubiquitin chains, in which the two proteins bind to distinct sites on NEMO. Moreover, the NZF1 domain simultaneously interacts with NEMO and Ile44 surface of a proximal ubiquitin from a linear diubiquitin chain, where the C-term tail of the ubiquitin is in the proximity of the NEMO ubiquitination site (Lys285). We further propose a model for the linear ubiquitination of NEMO by HOIP. In the model, NZF1 binds the monoubiquitinated NEMO and recruits the catalytic domains to the ubiquitination site, thereby ensuring site-specific ubiquitination of NEMO.

## Introduction

Nuclear factor-κB consists of a family of DNA transcription factors that regulate the expression of various genes involved in innate and adaptive immunity. In resting cells, the NF-κB proteins remain in the cytoplasm due to their interaction with inhibitor proteins known as inhibitors of κBs (iκBs) (Hoesel and Schmid, 2013; Park and Hong, 2016). NF-κB signaling is activated upon phosphorylation of the inhibitory molecules by the ikB kinase (IKK) complex through either canonical or non-canonical (alternative) pathways. In the canonical pathway, the IKK complex is composed of two catalytic (Ikkα and Ikkβ) and a regulatory subunit, IKKγ or NEMO (NF-κB essential modulator). Activation of the IKK complex and consequently the NF-κB pathway is contingent upon covalent and non-covalent interactions of NEMO with linear ubiquitin chains (Rahighi et al., 2009; Tokunaga et al., 2009). In linear ubiquitin chains, the C-terminal Gly of the first ubiquitin is attached to the Met-1 residue of the next ubiquitin in the chain.

Non-covalent interactions of NEMO with ubiquitin chains are mediated through its UBAN domain (ubiquitin-binding domain in ABINs and NEMO), encompassing amino acid residues 296-327 in human NEMO (Wagner et al., 2008). The UBAN domain forms a coiled-coil homo-dimeric structure, providing two highly symmetrical binding sites for linear diubiquitins (Lo et al., 2009; Rahighi et al., 2009). The two ubiquitin moieties in a linear diubiquitin chain use distinct surfaces for interacting with NEMO. While the N-terminal (distal) ubiquitin binds to the UBAN domain mainly via its hydrophobic Ile44 patch and C-terminal tail, the C-terminal (proximal) ubiquitin uses the polar residues adjacent to the Ile44 surface to interact with NEMO (Rahighi et al., 2009).

Covalent attachment of ubiquitin to NEMO (ubiquitination) occurs on Lys285 and Lys309 (human NEMO) and is catalyzed by the linear ubiquitin chain assembly complex (LUBAC). LUBAC is a multi-subunit E3 ubiquitin ligase complex composed of HOIP, HOIL-1L, and Sharpin, among which HOIP and HOIL-1L are RING-IBR-RING (RBR)-type E3 ligases (Ikeda et al., 2011; Tokunaga et al., 2009, 2011). HOIL-1L and Sharpin associate with HOIP through their ubiquitin-like (UBL) domains and relieve its autoinhibition (Liu et al., 2017).

HOIP is the main catalytic subunit of LUBAC. It contains a linear ubiquitin chain-determining domain (LDD), which along with RING2, specifies the formation of the linear (head-to-tail) type of ubiquitin chains (Liu et al., 2017; Smit et al., 2012; Stieglitz et al., 2013). While RING1 recognizes the ubiquitin-charged E2, the catalytic cysteine (C885) on RING2 forms a thioester intermediate with the donor ubiquitin. The LDD catalyzes the last step of the ubiquitin transfer to the substrate as it docks the acceptor ubiquitin, to which the donor ubiquitin is transferred (Smit et al., 2013; Stieglitz et al., 2013). Although the catalytic domain of HOIP is sufficient to synthesize free ubiquitin chains *in vitro*, the NZF1 domain of HOIP plays an important role in the process of linear ubiquitination through recognizing and binding the NEMO substrate (Fujita et al., 2014; Tokunaga et al., 2009). Mutations on the surface of NZF1 that binds to NEMO significantly reduce the level of linear ubiquitination of NEMO and subsequently NF-κB activation (Fujita et al., 2014). Moreover, HOIL-1L is reported to prime the linear ubiquitination process by ligating the first ubiquitin to the target lysine residue on NEMO (Kelsall et al., 2019; Smit et al., 2013). HOIL-1L is also suggested to regulate LUBAC activities by catalyzing mono-ubiquitination of the LUBAC components, followed by the attachment of linear ubiquitin chains catalyzed by HOIP (Fuseya et al., 2020).

Although LUBAC is identified as the E3 ligase responsible for linear ubiquitination of NEMO, detailed mechanistic information on how ubiquitin molecules are transferred to the NEMO substrate is still missing. Here, we report a crystal structure of NEMO in complex with the NZF1 domain of HOIP and linear diubiquitins. In this structure, HOIP NZF1 interacts with NEMO and ubiquitin simultaneously, suggesting that NZF1 recognizes the monoubiquitinated NEMO leading to the synthesis of linear ubiquitin chains by the catalytic domains of HOIP.

## Results

### NEMO CoZi binds HOIP NZF1 and two linear diubiquitin chains

While the NZF1 domain of HOIP is shown to bind NEMO (Fujita et al., 2014), it is still unclear how it cooperates with the catalytic domains for synthesizing linear ubiquitin chains. To further analyze the role of HOIP NZF1 in the linear ubiquitination of NEMO, we crystallized hNEMO (aa. 257-346) in complex with NZF1 (aa. 350-379) and linear diubiquitin chains (Fig.1A, B, Table 1). In the crystal structure, a NEMO dimer binds two linear diubiquitins on either side of the UBAN and a HOIP NZF1 at a region upstream of the UBAN domain. The NZF1-binding site is distinct and separated from the UBAN domain by ~25 Å on the NEMO coiled-coil axis (Fig. 1B). In this structure, the linear diubiquitin- and NZF1-binding surfaces on NEMO closely resemble the surfaces observed in the heterodimeric structures (PDB ID: 2ZVO, 2ZVN, and 4OWF) (Supplementary Fig. 1A). Despite minor differences in the orientation of linear diubiquitins with respect to the UBAN domain, NEMO/linear diubiquitin chains adopt similar binding modes to the previously reported structures that did not contain NZF1 (Fig. 1C, Supplementary Fig. 1B). In the crystal structure, the two linear diubiquitins bury surface areas of 1075.5 Å^2^ and 930.4 Å^2^ on NEMO. Each linear diubiquitin interacts with UBAN through the Ile44 hydrophobic patch as well as the C-terminal tail of the distal ubiquitin and Phe4-centered surface of the proximal ubiquitin (Fig. 1C). Such mode of interaction leaves the Ile44 patch of the proximal ubiquitins unoccupied and available for interactions with other binding partners.

**Table 1.**
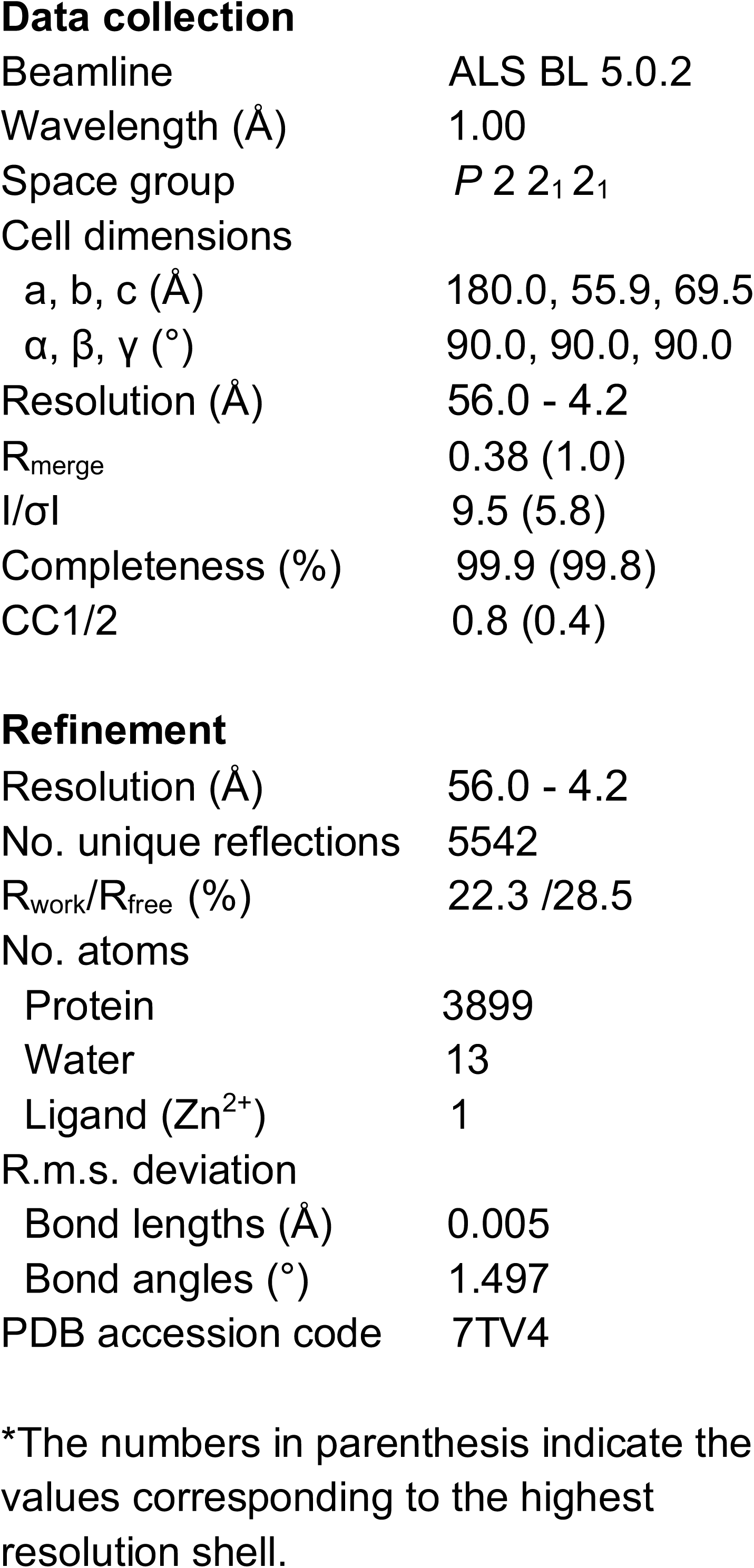
Data collection and refinement statistics

**Figure 1.**
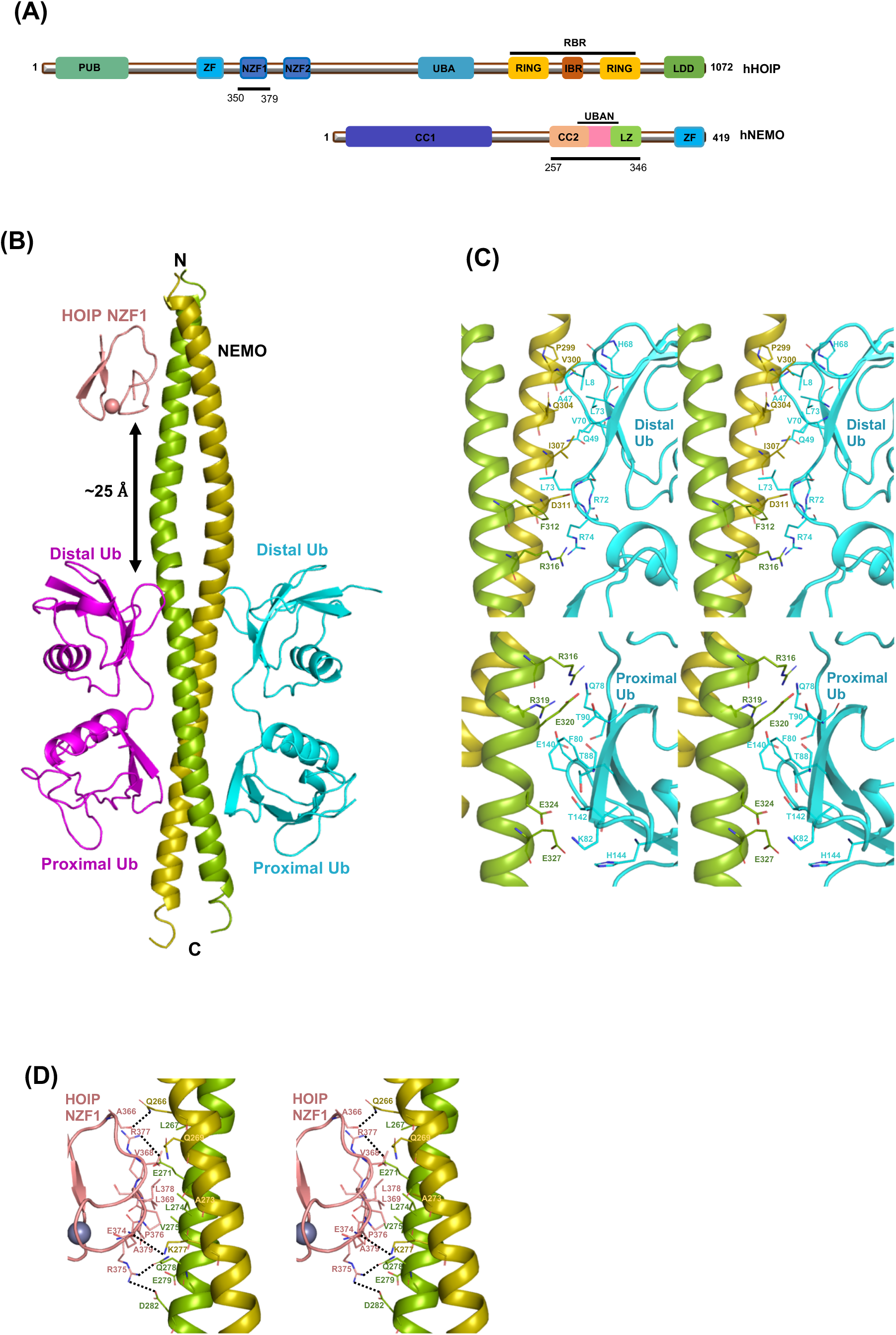
Crystal structure of NEMO CoZi in complex with HOIP NZF1 and linear diubiquitins. (A) Schematic domain composition of HOIP and NEMO proteins. The constructs of HOIP and NEMO used for crystallization are also indicated. PUB: PNGase/UBA or UBX-containing proteins, UBA: Ubiquitin-associated domain, UBAN: ubiquitin binding in ABIN proteins and NEMO, ZF: Zinc finger, NZF: Npl4 zinc finger, RING: Really interesting new gene, IBR: In between RING, RBR: RING-in-between-RING, LDD: Linear ubiquitin chain determining domain, CC: Coiled-coil region, LZ: Leucine zipper. (**B**) Overall structure of NEMO CoZi, HOIP NZF1 and linear diubiqutin heterotrimeric complex. The NZF1 and linear diubiquitin binding sites on NEMO are separated by 25 Å. The Stereo view of the interactions of NEMO with linear diubiquitin (**C**) and HOIP NZF1 (**D**).

The HOIP NZF1-binding site covers a surface area of ~516 Å^2^ on the NEMO dimer, involving amino acid residues ranging from Gln266 to Asp282 in the interactions with NZF1 (Fig. 1D). The NEMO-binding site on HOIP NZF1 is centered on a region located at the C-terminus of the Npl4 zinc finger, encompassing residues Glu374, Arg375, Pro376, Arg377. In addition to hydrophobic interactions, there are hydrogen bonds and salt bridges formed between Gln266/Ala366, Glu271/V368, Glu271/Arg377, and Lys277/Glu374, Gln278/Arg375, and D282/Arg375 of NEMO/HOIP NZF1 (Fig. 1D).

The two-fold symmetry of the NEMO dimer provides two potential binding sites on either side of the coiled-coil structure. However, unlike linear diubiquitin, only one NZF1 binds to NEMO in the heterotrimeric structure (2:2 stoichiometry for NEMO:diubiquitin vs. 2:1 stoichiometry for NEMO: NZF1). Such binding characteristic was also observed in the previously reported heterodimeric structure (Fujita et al., 2014) and can be attributed to the low affinity of NEMO for the isolated NZF1 domain, which requires increased local concentrations of the proteins for binding. Moreover, in the cellular context, the steric hindrance caused by the other domains in the full-length proteins may also lead to HOIP occupying one binding site on the NEMO dimer at a time.

### HOIP NZF1 interacts with NEMO and Ile44 surface of a proximal ubiquitin simultaneously

The Npl4 zinc fingers (NZF) are ubiquitin-binding domains (UBDs) that bind ubiquitin through a surface that is centered on three highly conserved amino acid residues T, F/Φ (Φ indicating a hydrophobic amino acid) (Alam et al., 2004; Wang et al., 2003). Interestingly, in the heterotrimeric crystal structure, the proximal ubiquitin from a linear diubiquitin of a symmetry-related molecule uses its Ile44 surface to interact with HOIP NZF1 (Fig 2A). The interactions of NZF1 with the proximal ubiquitin are primarily mediated through T360, F361/I372. The NZF domains in HOIP NZF1/proximal ubiquitin and Npl4 zinc finger/ubiquitin (PDB ID: 1Q5W) crystal structures adopt a highly similar binding mode for ubiquitin as demonstrated by a root mean square deviation (RMSD) of 1.3Å for the superimposition of Cα atoms of the NZF and ubiquitin molecules (Fig. 2B). Other residues from HOIP NZF1 that are engaged in interactions with ubiquitin include Ser358, Cys359, Gly362, and Ser371, among which backbone oxygen atoms of Ser358 and Ser 371 make hydrogen bonds with His68 and Arg42 of ubiquitin, respectively (Fig. 2C).

**Figure 2.**
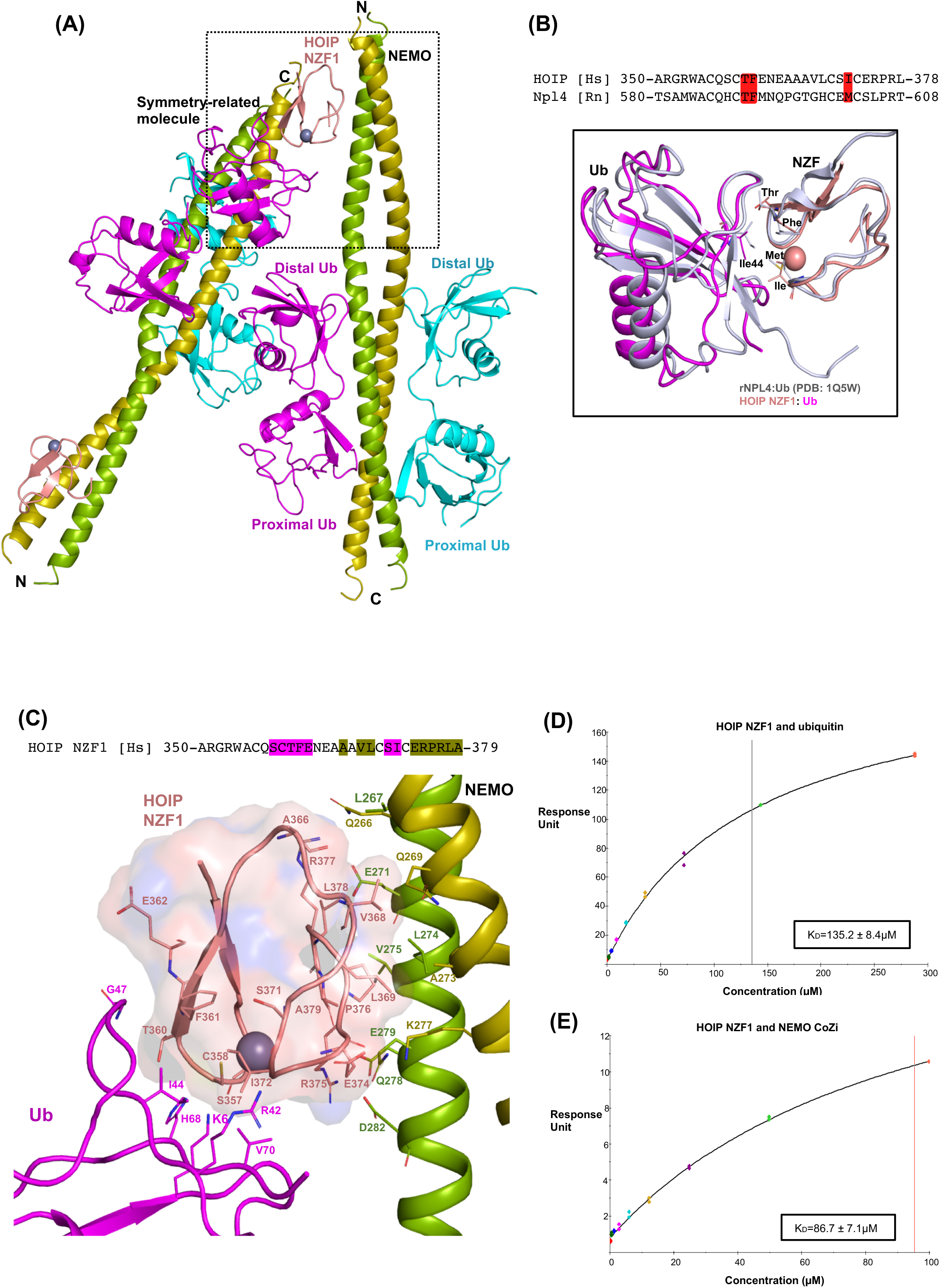
HOIP NZF1 binds NEMO and ubiquitin, simultaneously. (**A**) Crystal packing of the heterotrimeric structure indicates interaction of a proximal ubiquitin from a symmetry-related molecule with HOIP NZF1. (**B**) Sequence alignment and superimposition of HOIP NZF1 and Npl4 zinc finger. The conserved residues involved in interactions with ubiquitin are highlighted as red. (**C**) NZF1 uses distinct surfaces for binding NEMO and ubiquitin. Amino acid residues from HOIP NZF1 interacting with NEMO or ubiquitin are highlighted green and magenta, respectively. (**D**), (**E**) Surface plasmon resonance (SPR) analysis of HOIP NZF1 interaction with ubiquitin and NEMO, indicating equilibrium dissociation constant (K_D_) values of 135.2 ± 8.4 µM and 86.7 ± 7.1 µM (K_D_ ± SE), respectively.

Notably, the ubiquitin-binding site on NZF1 is distinct from the NEMO-binding site, and amino acid residues from HOIP that interact with ubiquitin and NEMO are not overlapping (Fig. 2C). Ubiquitin covers a surface area of 366.4 Å^2^ on the zinc-binding region of NZF1, which is slightly smaller than the NEMO-binding surface on NZF1 (516 Å^2^). Such small binding interfaces suggest a low binding affinity of HOIP NZF1 for either of the NEMO or ubiquitin molecules. To further characterize the binding of NZF1 with NEMO and ubiquitin, we performed surface plasmon resonance (SPR) measurements. The results demonstrate K_D_ of 86.7 µM and 135.2 µM for binding of HOIP NZF1 to NEMO and monoubiquitin, respectively (Fig. 2D, E). The low-affinity binding of HOIP NZF1 to either NEMO or ubiquitin may suggest a preference of HOIP for binding to the ubiquitinated NEMO over the individual proteins.

### A structural model of the recognition and linear ubiquitination of NEMO by HOIP

While the structural basis of free linear ubiquitin chain formation by HOIP RBR-LDD is described (Lechtenberg et al., 2016; Smit et al., 2012, 2013; Stieglitz et al., 2012, 2013), the mechanism through which NEMO is ubiquitinated by LUBAC is still unclear. Interestingly, in the NEMO/HOIP NZF1/linear diubiquitin complex crystal structure, the C-terminal tail of the proximal ubiquitin bound to HOIP NZF1 is in the proximity of NEMO Lys285, which is one of the two lysine residues reported as linear ubiquitination sites on human NEMO (Tokunaga et al., 2009a) (Fig. 3A). Notably, in the crystal structure, the last four residues at the C-terminus of the proximal ubiquitin (Leu73, Arg74, Gly75, and Gly76) are not visible in the electron density map. But the distance between the α-carboxyl of Arg72 and the sidechain ε-amine of Lys285 is ~12Å. However, considering the conformational flexibility of the ubiquitin tail, 12Å is sufficient for accommodating the remaining amino acids and forming an isopeptide bond with Ly285 of NEMO.

**Figure 3.**
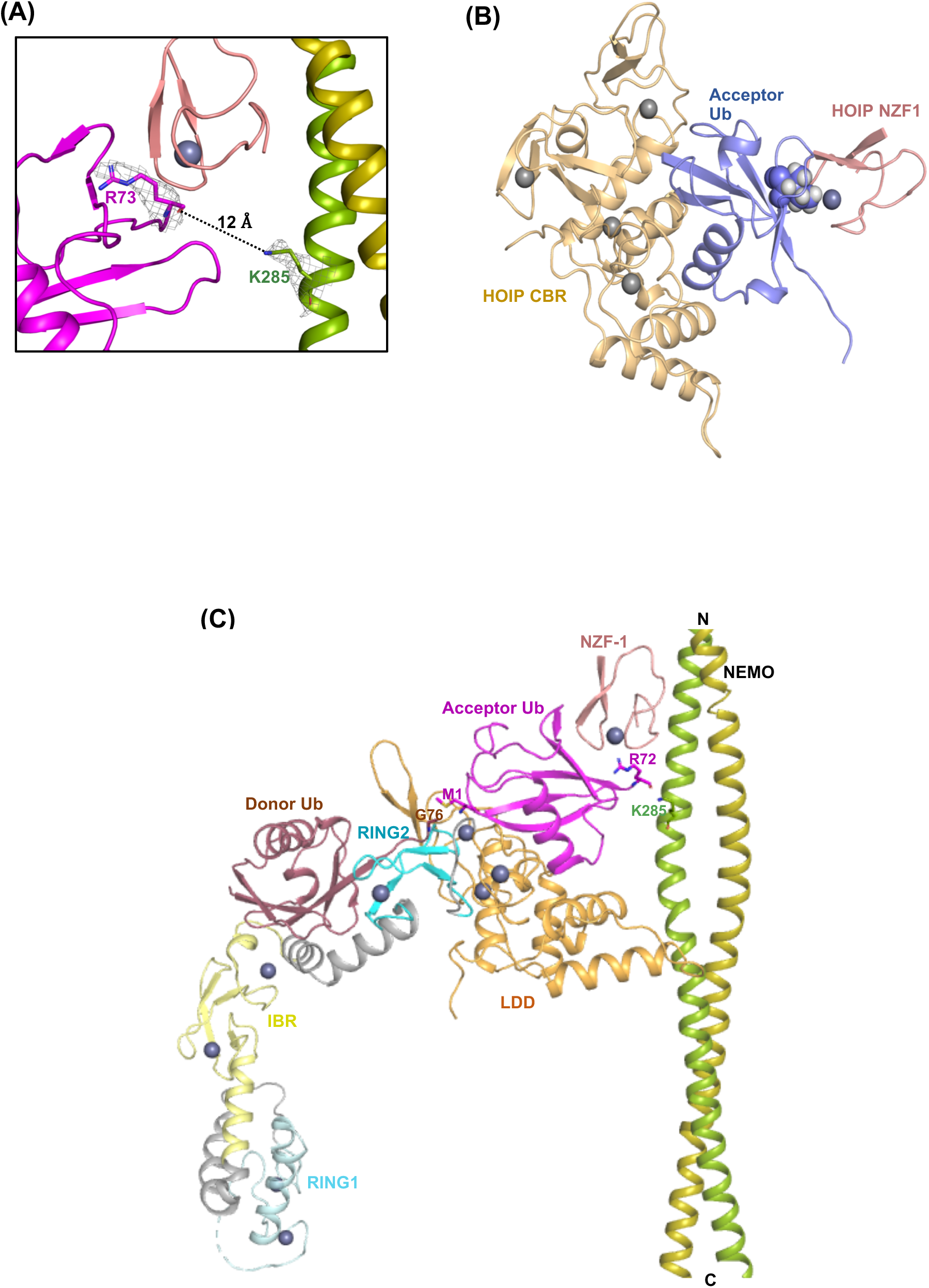
A structural model for the linear ubiquitination of NEMO by the HOIP E3 ligase. (**A**) The last four C-terminal amino acid residues are not visible in the electron density map of the heterotrimeric structure. The distance between the main chain carboxyl group of ubiquitin Arg72 and sidechain ε-amino group of NEMO Lys285 is ~12 Å, which is sufficient to accommodate the last four C-terminal residues of ubiquitin (a.a. 73-76) and form an isopeptide bond. The electron density indicate 2Fo-Fc map contoured at 1σ (**B**) Superimposition of the proximal ubiquitin on the acceptor ubiquitin bound to LDD (PDB ID: 4LJO), suggesting that ubiquitin can interact with NZF1 and RING2-LDD of HOIP. (**C**) Superimposition of the three complex structures including RING2-LDD/acceptor ubiquitin (PDB ID: 4LJO), RBR/donor ubiquitin (PDB ID: 5EDV), and NEMO/NZF1/ubiquitin (current structure) demonstrates a model for the site-specific ligation of linear ubiquitin chains to the NEMO substrate by HOIP.

On the other hand, while the conventional hydrophobic patch of the proximal ubiquitin is engaged in interactions with NZF1, the surface positioned at the opposite side from the Ile44 surface remains vacant and its orientation allows for the docking of the RING2-LDD of HOIP (Fig. 3B). Here, by consolidating the information obtained from the previously reported crystal structures of HOIP catalytic domains and the current crystal structure containing the NZF1, NEMO substrate, and ubiquitin we propose a model for the site-specific linear ubiquitination of NEMO by HOIP (Fig. 3C). In this model, the ubiquitin molecule in the proximity of NEMO ubiquitination site (K285) is sandwiched by the RING2-LDD and NZF1 domains, thereby suggesting that the proximal ubiquitin may be the acceptor ubiquitin for the linear ubiquitin chain synthesis. The model proposes a mechanism for cooperation of the various domains of HOIP in the ubiquitination process by ensuring (i) proper positioning of the N-terminal Met residue of the acceptor ubiquitin with respect to the HOIP active site and donor ubiquitin, and (ii) site-specific ligation of linear ubiquitin chains to NEMO.

## Discussion

The HOIP RBR-LDD is the main catalytic component of LUBAC involved in synthesizing free ubiquitin chains, but it is not sufficient for linear ubiquitination of substrates other than ubiquitin (Smit et al., 2013; Stieglitz et al., 2013). Also, HOIP does not initiate the substrate ubiquitination process. In this regard, it is suggested that HOIL-1L plays a role by priming linear ubiquitination through adding the first ubiquitin to the substrate followed by HOIP synthesizing and elongating the linear chain with a high processivity (Smit et al., 2013). *In vitro*, using isolated proteins, addition of catalytically active HOIL-1L to HOIP RBR-LDD significantly increases the ubiquitination levels of NEMO when compared with the RBR-LDD alone or in complex with a catalytically inactive mutant of HOIL-1L (C460A) (Smit et al., 2013). Although the NZF1 domain of HOIP has been shown to be responsible for the recognition of NEMO (Fujita et al., 2014; Tokunaga et al., 2009), it appears to be dispensable for NEMO ubiquitination when using isolated domains of NEMO and HOIP, *in vitro* (Smit et al., 2012). In the heterotrimeric structure, the concurrent interactions of NZF1 with ubiquitin and NEMO positions the C-terminus of ubiquitin in the proximity of the K285 ubiquitination site on NEMO. Superimposition of the proximal ubiquitin and the acceptor ubiquitin bound to CBR-LDD (PDB ID: 4LJO) indicates that the ubiquitin molecule interacting with both NEMO and NZF1 can be the acceptor ubiquitin bound to the RNG2-LDD of HOIP (Fig. 3B). The observation leads us to suggest that HOIP NZF1 recognizes the monoubiquitinated NEMO and guides the catalytic domains to the ubiquitination site to catalyze the synthesis of linear ubiquitin chains.

While there are two lysine residues in the CoZi domain of NEMO that are subjected to ubiquitination (Lys285 and Lys309 in human NEMO), it is yet unclear whether the two sites are ubiquitinated simultaneously or through alternative signaling cascades. Lys309 is part of the UBAN domain but it is not involved in interactions with diubiquitins (Fig. 1C). However, if Lys309 is ubiquitinated the ubiquitin chain may not bind the UBAN domain on the same NEMO molecule due to the steric hinderance. On the other hand, HOIP NZF1 does not appear to take part in the recognition and ubiquitination of NEMO Lys309 in the same way as Lys285, as its binding site on NEMO is ~40 Å away from Lys309. Therefore, we suggest that the presence of NZF1 facilitates the Lys285-specific ubiquitination of NEMO.

The heterotrimeric structure and the proposed model provide another piece of the puzzle of substrate ubiquitination by the LUBAC E3 ligase. However, still several other questions regarding the mechanism of substrate linear ubiquitination remain to be addressed. Although HOIL-1L is suggested to prime ubiquitination, its mechanism is yet to be explained. Moreover, the involvement of Sharpin in the linear ubiquitination process needs to be further investigated. Defining the roles of the two other subunits of LUBAC in the linear ubiquitination process will provide the rationale for LUBAC being evolved as a large multi-subunit E3 ligase complex.

## Acknowledgments

We would like to thank ALS BL-5.0.2 staff for their support of the diffraction data collection. This work was partially supported by the Chapman University Office of Research Faculty Opportunity Fund.

## Author contributions

Conceptualization, S.R; Methodology, S.R., Investigation, S.R., M.I., H.O., S.N, V.D.; Resources, S.R., Writing, S.R., M.I., H.O., Supervision, S.R.; Funding Acquisition, S.R.

## Declaration of interests

The authors declare no competing interests.

## Experimental Procedures

### Construction of plasmids

All plasmids were prepared for the bacterial expression system. Human NEMO (a.a. 257-346) and HOIP NZF1(a.a. 350-379) were cloned into pGEX-6p-1 vector and linear di-ubiquitin was cloned into pGEX-4T-1 for expression as N-terminally GST-tagged proteins. For SPR assays, mouse NEMO (a.a. 250-339) was cloned into a pET-28a vector downstream of the 6xHis tag.

### Protein expression and purification

Human NEMO and HOIP NZF1 plasmids were transformed into BL21 (DE3) E. coli cells (NEB) and expressed as GST-fusion proteins. The cells were grown in LB media containing 100 µg/mL ampicillin. Expression was induced by adding 0.25 mM IPTG once the OD600 was between 0.6 to 0.8. Proteins were expressed overnight at 25°C. The cells were harvested by centrifugation at 8000 rpm for 10 minutes at 4 °C and resuspended using phosphate-buffered saline (PBS). The cells were lysed on ice using a sonicator (QSonica). The lysate was cleared by centrifuging at 16000 rpm at 4°C. The supernatant was incubated with Glutathione Sepharose 4B (GS4B, Cytiva) for 2 hours on a rotating platform at 4°C. The beads were separated using a gravity column and washed multiple times using PBS to remove any unbound proteins. The GST tag was cleaved on column using thrombin protease for linear diubiquitin at 20°C overnight and PreScission protease for hNEMO and HOIP NZF1 at 4°C overnight. Cleavage was confirmed using SDS-PAGE. The cleaved proteins were eluted using PBS and further purified using size-exclusion chromatography on a Superdex 75 16/60 column (Cytiva). The running buffer used for size-exclusion contained 20 mM Tris-HCl, pH 8.0, and 150 mM NaCl. The pET28a-NEMO plasmid was transformed into BL21 (DE3) E. coli cells (NEB) and expressed as a His-tagged protein in LB media supplemented with 60 µg/mL of kanamycin. Expression was induced by the addition of 0.25 mM IPTG once the OD600 was between 0.6 to 0.8, at 25 °C overnight. The cells were collected and lysed as mentioned above. Once the lysate was cleared by centrifugation, the His-NEMO was separated using Talon beads (Cytiva). The 6xHis-tag was cleaved on column using thrombin protease at 20°C overnight. Cleavage was confirmed using SDS-PAGE. NEMO was eluted using PBS and purified using size-exclusion chromatography as described above.

### Surface Plasmon Resonance (SPR)

The SPR assays were performed using a Biacore S200 (Cytiva). Anti-GST(α-GST) antibodies were covalently immobilized on the carboxymethyl dextran (CM5) sensor chip by amine coupling. The surface of the CM5 chip was first activated using EDC/NHS followed by immobilization of the α-GST antibodies and deactivation using ethanolamine. GST (control) and GST-HOIP NZF-1(a.a. 350-379) were immobilized on the sensor chip containing α-GST antibodies. Various concentrations of mono-ubiquitin and mNEMO (a.a. 250-339) were prepared in the running buffer containing 10 mM HEPES, pH 7.4, 150 mM NaCl, and 0.005% P20. Each experiment was performed in duplicate.

### Crystallization, x-ray diffraction data collection, and structure determination

The hHOIP NZF1 (a.a. 350-379), hNEMO (a.a. 257-346), and linear di-ubiquitin proteins were purified separately and mixed in an equimolar ratio for crystallization. The co-crystals of the three proteins were obtained in sitting drops containing 0.1 M Tris-HCl (pH 8.5), and 22 % v/v PEG Smear Broad after six weeks. X-ray diffraction data was collected at the Advanced Light Source Beamline 5.0.2 (ALS BL-5.0.2) at 100 and processed using iMosflm. The structure was solved by molecular replacement in MOLREP (Vagin and Teplyakov, 2010) using structures of NEMO/HOIP NZF1 (PDB ID: 4OWF) and ubiquitin (PDB ID: 1UBQ) as search models. The model was further built in COOT (Emsley and Cowtan, 2004)and refined using REFMAC5 (Murshudov et al., 2011). Data collection and refinement statistics are summarized in Table 1. Structural figures are made using PyMOL (PyMol Molecular Graphics System, Version 2.2.3; Schrodinger).

### Accession numbers

Atomic coordinates and structure factors of the NEMO CoZi in complex with HOIP NZF1 and linear diubiquitins is deposited to the protein data bank (PDB) with accession code 7TV4.

## Notes

### Competing Interest Statement

The authors have declared no competing interest.

