## Supplemental Figure 1 for "Structural basis for the simultaneous recognition of NEMO and acceptor ubiquitin by the HOIP NZF1 domain"

**Supplementary Figure 1. (related to Fig. 1)**

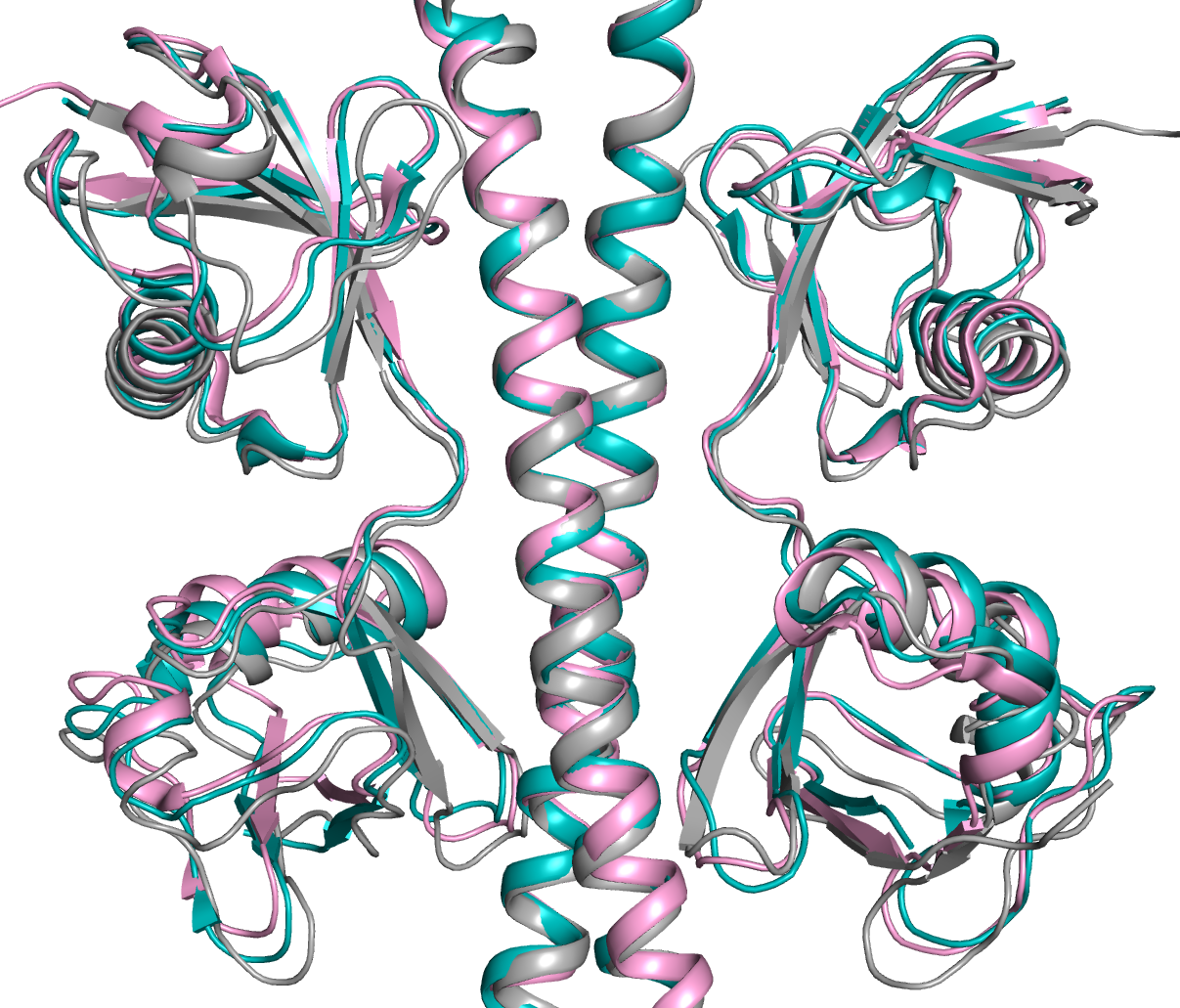

**Distal Ub**

**Distal Ub**

**Proximal Ub**

**Proximal Ub**

**NEMO**

**2ZVO**

**2ZVN**

**Current structure**

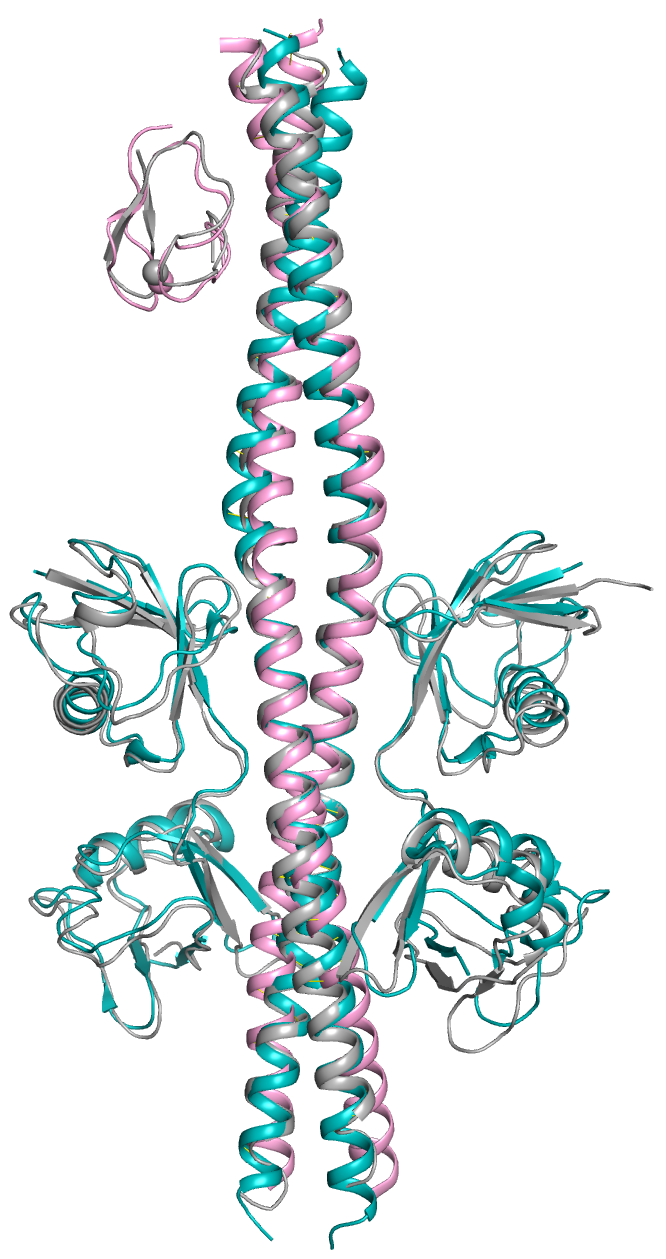

**N**

**C**

**Distal Ub**

**Proximal Ub**

**Distal Ub**

**Proximal Ub**

**NEMO**

**HOIP NZF1**

**(B)**

**4OWF**

**2ZVN**

**Current structure**

**(A)**

**Supplementary Figure 1.** NEMO in the heterotrimeric structure adopts the same binding mode for HOIP-NZF1 and linear diubiquitins as the heterodimeric structres.

(A) Superimposition of the C⍺ atoms of NEMO CoZi in the heterotrimeric crystal structure (Grey) on NEMO in heterodimerc complex structures containing HOIP NZF1 (PDB ID: 4OWF, pink) and diubiquitions (PDB ID: 2ZVN, cyan). (B) Superimposition of the C⍺ atoms of UBAN in the heterotrimeric structure (Grey) on the UBAN in crystal structures containing linear diubiquitins (2ZVO and 2ZVN).
